# Structural analysis of the master regulator Rns reveals a small molecule inhibitor of enterotoxigenic *Escherichia coli* virulence

**DOI:** 10.1101/2020.10.05.326769

**Authors:** Charles R Midgett, Kacey Marie Talbot, Jessica L. Day, George P Munson, F Jon Kull

## Abstract

Enterotoxigenic *Escherichia coli* (ETEC) is a common cause of diarrheal disease worldwide and a frequent cause of travelers’ diarrhea. In addition to the production of enterotoxins, studies with human volunteers established ETEC virulence is dependent upon the production of proteinaceous adhesive pili for attaching to the intestinal wall. Although pilins are highly immunogenic, vaccines incorporating them have yet to be proven efficacious. An additional challenge for vaccines is the heterogeneity of ETEC pili, as 20 different pilus types have been identified. However, the expression of a significant number of pilus types is dependent upon Rns, an AraC family transcription factor. Furthermore, Rns also regulates the expression of the virulence factor CexE, an outer membrane coat protein. To determine how Rns functions and is regulated we solved its structure by X-ray crystallography to 3 Å resolution. Rns forms a dimer via its N-terminal domain and its structure is consistent with the dimer binding looped DNA. Our analyses also revealed a fatty acid, decanoic acid, bound within the Rns structure. Although Rns was not known to specifically bind small molecule ligands, biochemical analysis showed decanoic acid specifically stabilized Rns in a dose dependent manner. Lac reporter assays further showed that decanoic acid inhibits Rns function at both activated and repressed promoters. In situ, exogenous decanoic acid inhibited the expression of Rns-dependent CFA/I pili and CexE in different ETEC strains. Thus, our study reveals for the first time a naturally occurring small molecule ligand specifically inhibits Rns activity and potently suppresses the expression of ETEC virulence factors. Our findings provide an alternative approach to vaccines for inhibiting ETEC pathogenesis by using the Rns structure as a framework for rational drug design.

## Introduction

Diarrheal diseases are estimated to cause nearly 1.3 million deaths every year (Troeger *et al*., 2017). Enterotoxigenic *E. coli* (ETEC) is a leading cause of diarrhea, however there are very different estimates of its prevalence. Older estimates suggested 280 million global infections with 380,000 deaths per year of children under five (World Health Organization, 1999), while newer estimates suggest only 75 million cases each year, killing up to 30,659 children under the age of five (Khalil *et al*., 2018). While there is debate about the worldwide incidence of ETEC it is clearly a leading cause of traveler’s diarrhea and is of particular concern during military deployments (Jiang and DuPont, 2017; Porter *et al*., 2017).

ETEC colonization is dependent on production of pili, sometimes referred to as colonization factors, that are required for the bacteria to attach to the intestinal wall. In the absence of adhesive pili, ETEC is unable to cause infection (Satterwhite *et al*., 1978; Turner *et al*., 2006). Over 20 of these highly immunogenic colonization factors have been identified (Fleckenstein and Kuhlmann, 2019), and while it has been hypothesized vaccines incorporating them could be protective from ETEC infection (Boedeker, 2005; Vidal *et al*., 2019), to date such vaccines have yet to demonstrate protective immunity (Boedeker, 2005; Qadri *et al*., 2020). This coupled with the rise of antibiotic resistant strains, with their associated treatment challenges and increased cost (Shakoor *et al*., 2019), shows it is critical to change vaccine strategy, as well as identify new avenues for treatment.

While ETEC colonization factors are diverse, nearly half are regulated by the transcription factor Rns (also known as CfaD or CfaR), a member of the AraC family (Caron *et al*., 1989; Caron *et al*., 1990; Bodero *et al*., 2008; Munson, 2013). Rns activates pilin promoters by binding to a proximal site immediately adjacent to the −35 hexamer (Munson and Scott, 1999; Munson and Scott, 2000; Bodero *et al*., 2007), which may be accompanied by one or more additional sites further upstream. Like most other AraC family members, Rns contains two helix-turn-helix DNA binding motifs in its carboxy terminal domain (Gallegos *et al*., 1997), as well as an amino terminal domain that has been suggested to be involved in dimerization (Mahon *et al*., 2010) and ligand binding (Gallegos *et al*., 1997). Notably, prior to this study ligand binding by Rns had not been experimentally demonstrated.

ToxT, an AraC family member from *V. cholerae* regulates transcription of the two major virulence factors, the toxin-coregulated pilus (TCP) and cholera toxin (CT) (DiRita *et al*., 1991; Higgins *et al*., 1992). Previous work in our laboratory identified unsaturated fatty acids (UFAs) inhibit ToxT activity through binding a pocket in the N-terminal domain. This inhibition is potentially due to a disruption in the ability of ToxT to bind to promoter sites as the presence of the monounsaturated fatty acids oleic (C18) and palmitoelic (C16) acids inhibited ToxT DNA binding *in vitro* (Lowden *et al*., 2010). As fatty acids are a component of bile, which is present in the GI tract of most organisms affected by *V. cholerae*, it is reasonable that such compounds would make an effective signal for virulence regulation (Chatterjee *et al*., 2007) and suggests other virulence controlling AraC family members might be inhibited by a similar mechanism (Lowden *et al*., 2010). A computational screen of AraC proteins identified several candidates for such regulation, including Rns.

In order to determine whether Rns is inhibited by UFA’s in a manner similar to ToxT, we solved the crystal structure of Rns. We observed two protein conformations, “open” and “closed” and show that Rns forms a dimer, consistent with it binding DNA as a dimer in a manner similar to other AraC proteins (Soisson *et al*., 1997; LaRonde-LeBlanc and Wolberger, 2000; Shrestha *et al*., 2015). Inspection of the open conformation revealed an unknown ligand bound in a groove between the N- and C-terminal domains, which was modeled as decanoic acid. Differential scanning fluorometry revealed decanoic acid increased the melting temperature (*T*_m_) of Rns, stabilizing it and supporting a model in which decanoic acid specifically binds to the protein. When Rns was crystallized with excess decanoic acid only the closed conformation was observed. In support of a physiological role for a fatty acid ligand, exogenous decanoic acid abolished the expression of colonization factors by inhibiting Rns-dependent transcriptional regulation. These results support the hypothesis that Rns, like ToxT, utilizes a common mechanism of binding fatty acid effector molecules to regulate virulence gene expression.

## Materials and Methods

### Identification of candidate AraC proteins

Fifty-eight proteins that had been previously characterized and identified as AraC family members were chosen for analysis (Ibarra *et al*., 2008). Although these proteins were linked to various functions (categories included: general metabolism, adaptive responses to nutrient sources, stress, and virulence), we analyzed all 58, regardless of functional class. As the prior study utilized sequence alignments for their analysis (Ibarra *et al*., 2008), we used alignments of the predicted secondary structure for our analysis. The following criteria were used to identify possible candidates: 1) The protein length was similar to that of ToxT. 2) The predicted secondary structure indicated a protein with an N-terminal β-strand rich domain and a C-terminal DNA binding domain comprised of seven α-helices (the canonical AraC DNA binding domain). 3) The presence of a positively charged amino acid at the C-terminal end of the α-helix predicted to be analogous to Lys230 in α7 of ToxT. 4) The presence of a positively charged amino acid in an appropriate position in the β-strand N-terminal domain. As β-strands are much more difficult to predict than α-helices, and as β-strand domains are inherently more variable, the exact location of this residue was more flexible than in rule #3. Using these rules, we identified four AraC virulence regulators out of the 58 that met all four primary criteria and for which comparison with an existing phylogenic tree (Ibarra *et al*., 2008) showed they were among the closest relatives of ToxT. From members of the initial 58 candidates that failed our virtual screen based on rule #3, we identified three additional AraC proteins that are involved in virulence and have a positively charged residue near the end of the helix (Fig. 1, Table 1).

**Table 1:** List of potential ToxT like AraC proteins

| **Table 1: List of potential ToxT like AraC proteins** |  |  |  |
| --- | --- | --- | --- |
| **Protein** | **Organism** | **UniProt ID** | **Criteria met** |
| FapR | *E. coli* – ETEC | P23774 | 1-4 |
| PerA | *E. coli* – EPEC | P43459 | 1-4 |
| SirC | *S. typhi* | Q8Z4A6 | 1-4 |
| HilD | *S. typhimurium* | P0CL08 | 1-4 |
| VirF | *S. flexneri* | P0A2T1 | 1, 2, 4 |
| VirF | *Y. enterocolitica* | P0C2V5 | 1, 2, 4 |
| Rns | *E. coli* – ETEC | P16114 | 1, 2, 4 |

**Figure 1:**
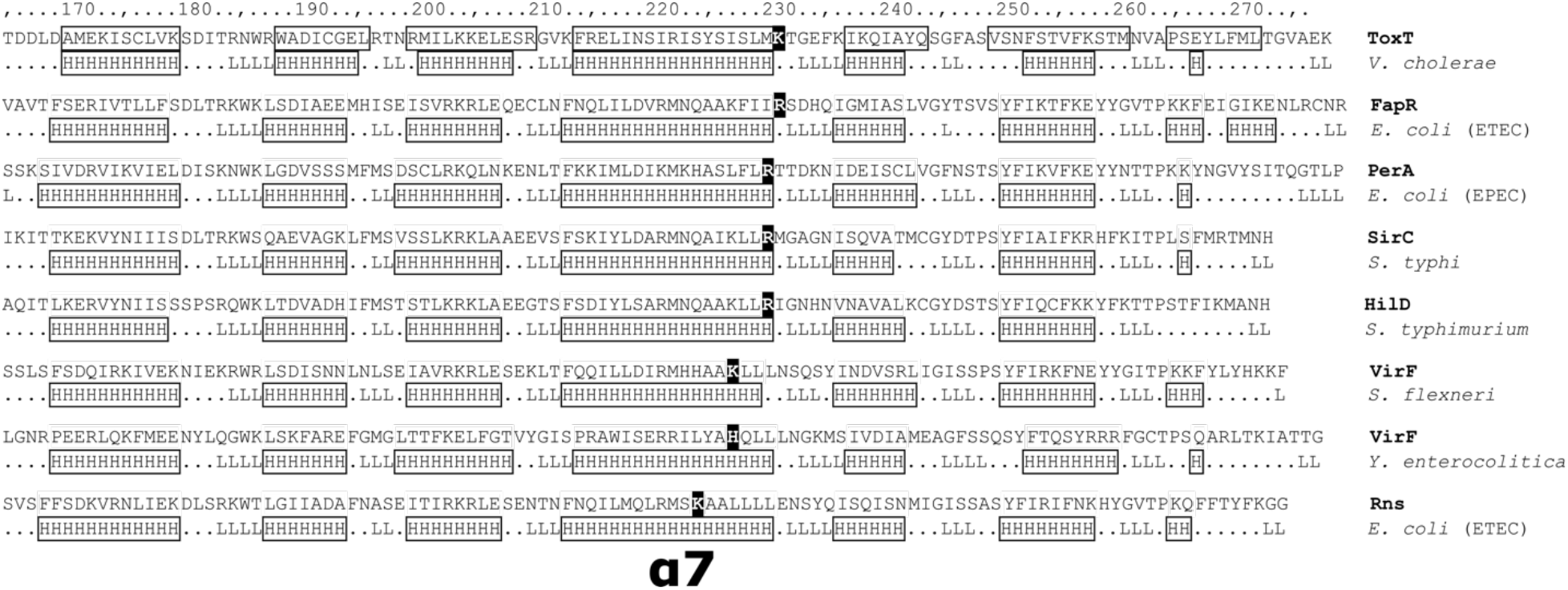
Structural alignment of the C-terminal domains of the top AraC family hits. ToxT is shown at the top with structurally determined α-helices shown in boxes. For the other proteins, H indicates predicted α-helices and L indicates predicted loop regions. The position of α7 is indicated and the positive residues are highlighted for each sequence. Predictions by the PROF algorithm using the server at http://predictprotein.org.

### Cloning of Rns into an expression vector

The sequence encoding Rns was optimized to remove rare codons and flanking sequences to insert into the plasmid pCDB24, a gift from Dr. Bahl (Institute for Protein Innovation), which contains a N-terminal 10xHis-SUMO tag (pCDB24 addgene.org), were added. The resulting construct was synthesized by IDT. The construct sequence was amplified using PCR and the resulting product was purified using the QIAquick PCR purification kit from Qiagen. The vector was digested with XhoI and purified using the QIAqucik PCR purification kit. The *rns* construct was cloned into the digested vector in frame with the SUMO tag, using the DNA Assembly Mix from NEB according to the manufacturer’s instructions, to create the 10xHis-SUMO-Rns (SMT-Rns) construct. The correct insertion of *rns* was verified by sequencing. The resulting plasmid was transformed into BL21 DE3 cells for unlabeled protein expression and into B834 DE3 cells for SeMet labeling.

### SeMet labeled protein expression

SeMet labeled protein was expressed using Selenomet media from Molecular Dimensions. The cultures were initiated from a frozen stock into Selenomet media supplemented with 50 *µ*g/ml of L-methionine and 100 µg/ml carbenicillin. The culture was incubated at 37 °C overnight with shaking. The next morning the culture was centrifuged at 25 °C for 10 mins at 3000 xg to collect the cells. The cells were washed three times with water. After washing the cells were resuspended in a 10 fold larger volume of Selenomet media supplemented with 50 µg/ml of seleno-L-methionine, and 50 µg/ml of carbenicillin. The culture was incubated at 37 °C till an OD600 of 1-2. The culture was induced by adding 500 µM IPTG and incubated at 18 °C overnight.

### Unlabeled Rns expression

SMT-Rns was expressed in modified TB media (Fisher Scientific). To start a frozen stock was used to inoculate a starter culture of 2 ml ZYP-0.8G media (ZYP media (Studier, 2005) supplemented with 0.8% glucose) with 200 µg/ml carbenicillin, which was incubated overnight at 30 °C. The next morning of the starter was diluted 1:100 in TB containing 2 mM MgSO_4_ and 200 µg/ml of carbenicillin. This culture was incubated at 37 °C till an OD600 of 2-3. The culture was then diluted 1:10 in TB with 2 mM MgSO_4_ and 50 µg/ml carbenicillin in a 3 L baffled flask. This was again incubated at 37 °C till it reached an OD600 of 2-3. The culture was then induced with 500 µM IPTG and 5% glycerol was added. The culture was incubated at 18 °C overnight.

### Rns purification

SMT-Rns purification was performed as follows. The culture was pelleted at 3000 xg for 25 mins. The bacteria were resuspended in wash buffer (20 mM TRIS pH 8, 20 mM imidazole, 500 mM NaCl) supplemented with 500 µM EDTA, 500 µM PMSF, and a Roche protease inhibitor tablet. The culture was lysed by three passes through a French press. The lysate was clarified by ultracentrifugation at ∼100,000 x g for 45 mins. The supernatant was filtered using a 0.45 µm filter, and 1 M MgCl_2_ was added to a final concentration of 1 mM.

The SMT-Rns was captured using a HisTrap column from GE Healthcare. The column was equilibrated with 10 CV of elution buffer (20 mM TRIS pH 8, 500 mM imidazole, 500 mM NaCl) followed by 10 CV of wash buffer. The supernatant was loaded onto the column at 2 ml/min. The column was washed with 10 CV of wash buffer followed by 9 CV of 9% elution buffer, and 2 CV of 20% elution buffer. SMT-RNS was eluted from the column with a 10 CV gradient of 20%-100% elution buffer. Fractions were collected in tubes with EDTA for a final concentration of 100 *µ*M.

Cleavage of the SMT-Rns was achieved using previously purified 6xHis-Ulp1-6xHis SUMO protease, a gift from Dr. Bahl (Institute for Protein Innovation). The relevant fractions from the His-Trap column were pooled and dialyzed against 2 L of (20 mM TRIS pH 8, 200 mM NaCl, 1 mM DTT, 500 *µ*M EDTA) at 4 °C. After ∼4 hours the dialysis buffer was changed, and a 10 mg aliquot of the protease was added to the pooled fractions. Dialysis was continued overnight at 4 °C. The next morning a HisTrap column was equilibrated with 5 CV of dialysis buffer with 1 mM MgCl_2_ and 20 mM imidazole. Imidazole was added to the dialysate for a final concentration of 20 mM and MgCl_2_ was added for a final concentration of 1 mM. The dialysate was then applied to the column and the flow through was collected.

Final purification was carried using an HiTrap Sp ion exchange column (GE Healthcare). The column was equilibrated with 10 CV of Sp elution buffer (20 mM TRIS pH 8, 1 M NaCl, 250 µM EDTA) followed by 10 CV of Sp wash buffer (20 mM TRIS pH 8, 200 mM NaCl, 250 µM EDTA). Then the flow through from the cleavage step was loaded onto the Sp column. The column was washed with 10 CV of Sp wash buffer. Elution was performed using a gradient to 100% Sp elution buffer over 5 CV. The concentrations of the relevant fractions were determined, and DTT to a final concentration of 1 mM was added to the fractions.

### Crystallization and data collection

Initial crystal screening for the SeMet labeled Rns was done using 96 well block screens from either Qiagen or Hampton Research. Initial drops were setup either at 2 mg/ml or 0.6 mg/ml using a NT8 robot. The crystals were imaged using Rock Imager. The initial hits were then optimized. The best crystals of Rns were obtained in a base condition of 0.1 M (D/L) malic acid pH 7, with 6%-10% PEG 3350. These conditions were then used in additive screens, Additive Screen (Hampton Research). The final crystallization condition for SeMet labeled Rns was.6 mg/ml protein added 1:1 to 0.1 M (D/L) malic acid pH 7, 6% PEG 3350, 0.01 M Betaine hydrochloride in sitting drops. The crystals were frozen using the well solution supplemented with 10% PEG 3350, and 40% glycerol as the cyro-protectant. Data was collected using the FMX beam line at NSLS-II. Two anomalous data sets were collected from the same crystal with 360° of rotation at a wavelength of 0.979184 Å.

Another set of screens were setup with native Rns at.6 mg/ml with 1 mM decanoic acid. The native Rns, decanoic acid mix was crystallized by adding the mixture in a 1:1 ratio to 0.1 M succinnic acid, 14% PEG 3350, 0.03 M glycyl-glycyl-gylcine in hanging drops. The crystals were frozen with the well solution supplemented with 30% ethylene glycol as the cryo-protectant. Diffraction data was collected at the AMX beam line at NSLS-II.

### Anomalous data processing and refinement

The anomalous data sets were processed using XDS (Kabsch, 2010). The space group was determined to be P 2_1_ with a unit cell of 72.51, 49.97, 102.92, 90.00, 106.11, 90.00. The two datasets were combined in XSCALE (Kabsch, 2010). The processed data was cut at 2.8 Å and used in a Hybrid Substructure Search followed by one round of AutoSol and one round of AutoBuild in PHENIX (Adams *et al*., 2010). The model was refined using iterative rounds of automated refinement with Refine as implemented in PHENIX with NCS, secondary structure, as well as experimental phase restraints (Adams *et al*., 2010), followed by manual model building using COOT (Emsley and Cowtan, 2004). Chimera and Chimerax were used for model visualization (Pettersen *et al*., 2004; Goddard *et al*., 2017). Refinement statistics are listed in Table 2.

**Table 2:** Refinement Statistics

| Table 2: Refinement Statistics |  |  |
| --- | --- | --- |
|  | SeMet-Rns | Rns |
| Wavelength |  |  |
| Resolution range | 29.29 - 2.8 (2.9 - 2.8) | 27.39 - 3.0 (3.107 - 3.0) |
| Space group | P 21 | P 21 21 21 |
| Unit cell | 72.51 49.97 102.92<br>90 106.11 90 | 48.24 95.15 133.09<br>90 90 90 |
| Total reflections | 228834 (24784) | 81535 (8041) |
| Unique reflections | 17538 (1743) | 12798 (1219) |
| Multiplicity | 13.0 (14.2) | 6.4 (6.6) |
| Completeness (%) | 94.93 (86.50) | 99.16 (97.50) |
| Mean I/sigma(I) | 8.67 (0.74) | 7.29 (0.88) |
| Wilson B-factor | 89.24 | 93.97 |
| R-merge | 0.1984 (3.508) | 0.1518 (1.994) |
| R-meas | 0.2069 (3.637) | 0.1654 (2.162) |
| R-pim | 0.0579 (0.956) | 0.06498 (0.8261) |
| CC1/2 | 0.995 (0.29) | 0.997 (0.549) |
| CC* | 0.999 (0.671) | 0.999 (0.842) |
| Reflections used in refinement | 16917 (1532) | 12745 (1211) |
| Reflections used for R-free | 1679 (143) | 1277 (120) |
| R-work | 0.2422 (0.3928) | 0.2571 (0.3840) |
| R-free | 0.2999 (0.4694) | 0.3007 (0.4249) |
| CC(work) | 0.948 (0.589) | 0.955 (0.646) |
| CC(free) | 0.911 (0.461) | 0.915 (0.423) |
| Number of non-hydrogen<br>atoms | 4222 | 4068 |
| macromolecules | 4189 | 4044 |
| ligands | 33 | 24 |
| Protein residues | 513 | 494 |
| RMS(bonds) | 0.008 | 0.012 |
| RMS(angles) | 1.17 | 1.36 |
| Ramachandran favored (%) | 93.89 | 92.39 |
| Ramachandran allowed (%) | 5.92 | 7.2 |
| Ramachandran outliers (%) | 0.2 | 0.41 |
| Rotamer outliers (%) | 4.99 | 6.9 |
| Clashscore | 7.4 | 19.06 |
| Average B-factor | 93.97 | 83.91 |
| macromolecules | 94.01 | 84.04 |
| ligands | 89.13 | 61.91 |

### Native data processing and refinement

The native data sets were processed using XDS (Kabsch, 2010). The space group was P2_1_2_1_2_1_ with a unit cell of 48.24 95.15 133.09 90 90 90. This crystal had significant pseudo-translational symmetry with an off-origin peak 46% of the origin peak as determined by PHENIX (Adams *et al*., 2010). The structure was solved using Phaser (McCoy *et al*., 2007) as implemented in PHENIX (Adams *et al*., 2010) using the previously solved SeMet-Rns as a search model. After phaser there was visible density in the ligand binding pockets of both monomers. The density was still visible after a cycle of automated refinement in PHENIX (Adams *et al*., 2010). Decanoic acid was added to the density using COOT (Emsley and Cowtan, 2004) and refinement was performed using iterative rounds of PHENIX Refine (Adams *et al*., 2010) followed by manual model building in COOT (Emsley and Cowtan, 2004). Chimera and Chimerax were used for model visualization (Pettersen *et al*., 2004; Goddard *et al*., 2017). Refinement statistics are listed in Table 2.

### Differential scanning fluorometry (DSF)

DSF was performed to assess the effect of fatty acids on Rns stability (Niesen *et al*., 2007). First stocks of fatty acids at 100x of final concentration were made by adding octanoic, decanoic, and palmitic acid to methanol. The decanoic acid was serially diluted by 1/2, in methanol, to obtain the concentrations for the dose response. 1 µl of the appropriate fatty acid was added to 99 µl of Rns, at a concentration of ∼.7 mg/ml, and incubated at room temperature for 1 hour. Then 18 µl of the mixture was added to a PCR plate in triplicate. Sypro Orange dye (Life Technologies), diluted in buffer, was added to the PCR plate for a final concentration 5x, and a total reaction volume of 20 *µ*l. For each condition a buffer only control was also performed in triplicate. The melting curves were generated using a StepOne + RT PCR machine (Life Technologies) with a gradient from 25 °C to 95 °C utilizing 1 °C steps. The normalized melt data was exported into STATA for analysis as described (Midgett *et al*., 2017).

### Plasmids

Plasmid pGPMRns-Myc is a derivative of pTags2 (Addgene) that expresses Rns-myc from *lacp. rns* was amplified from pGPMRns (Bodero *et al*., 2007) with primers 1519/1522. pTags2 vector backbone was amplified with primers 1414/1419. The two PCR products were Dpn1 digested then circularized with NEB HiFi. All plasmids used in this study are listed on table 3 and oligo-nucleotides are listed on table 4.

**Table 3:**
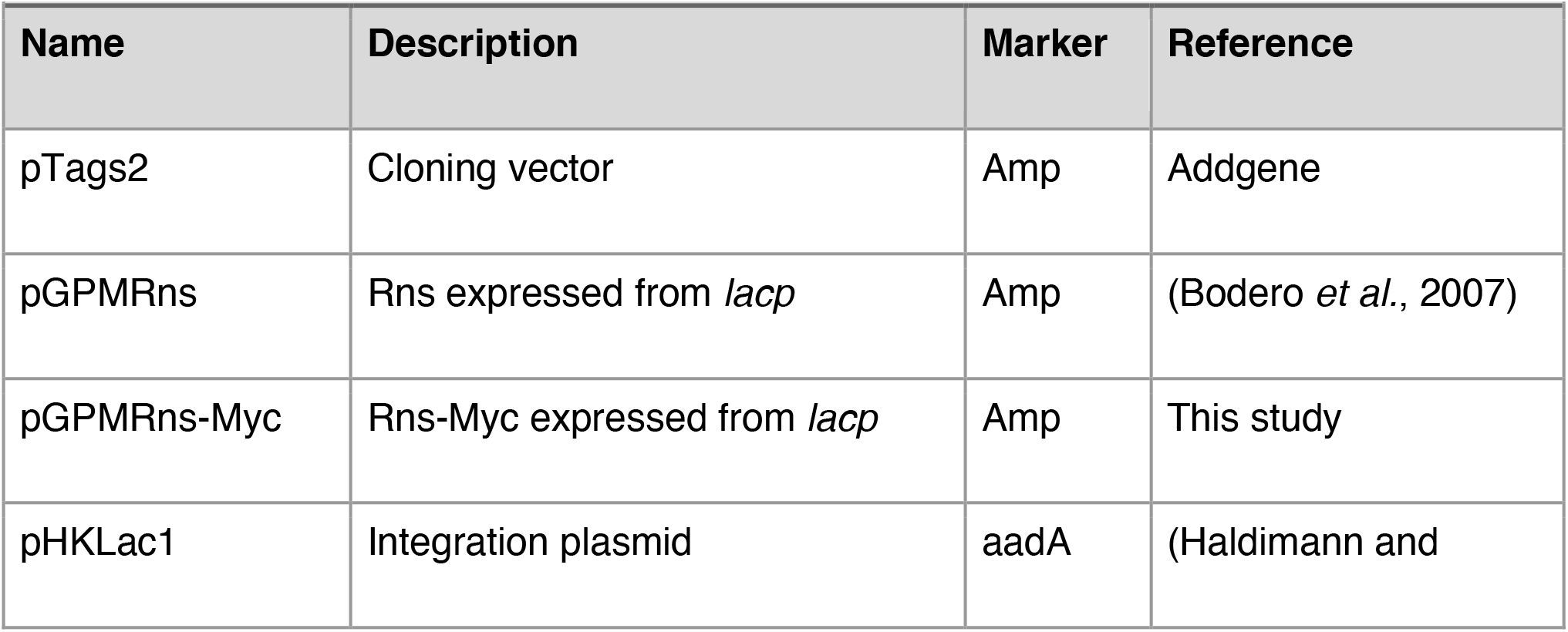
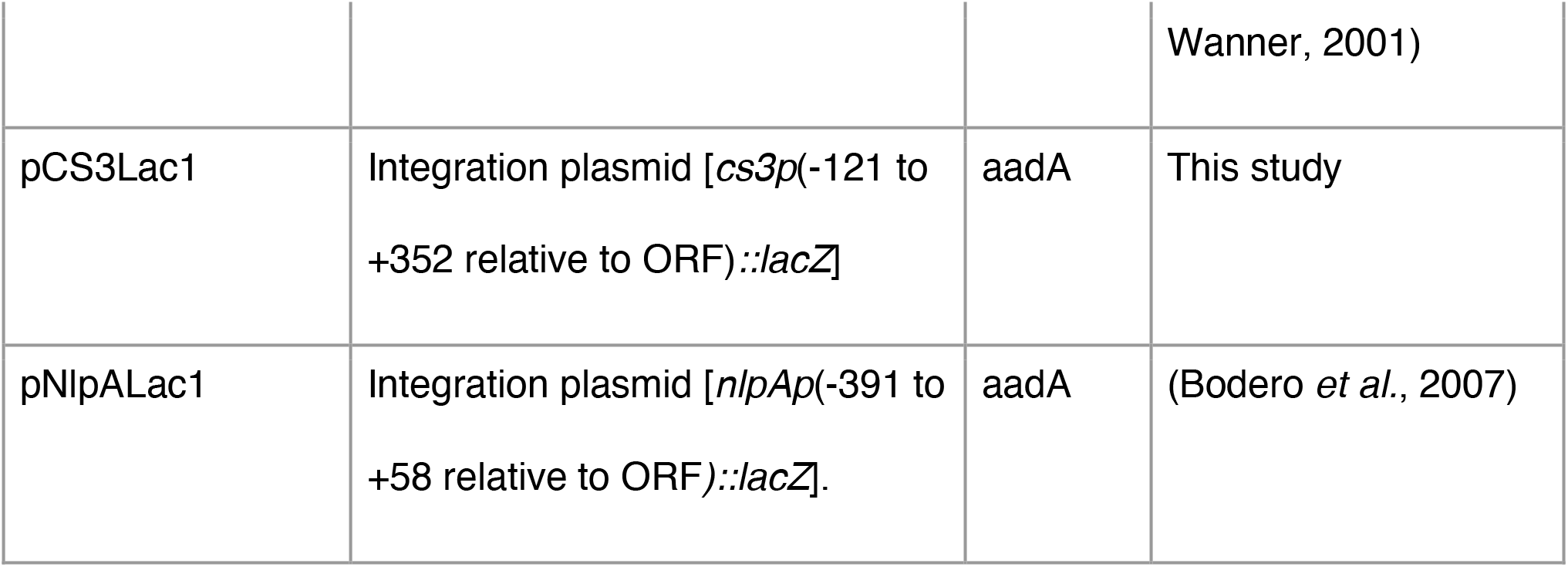
Plasmids used in this study

**Table 4:**
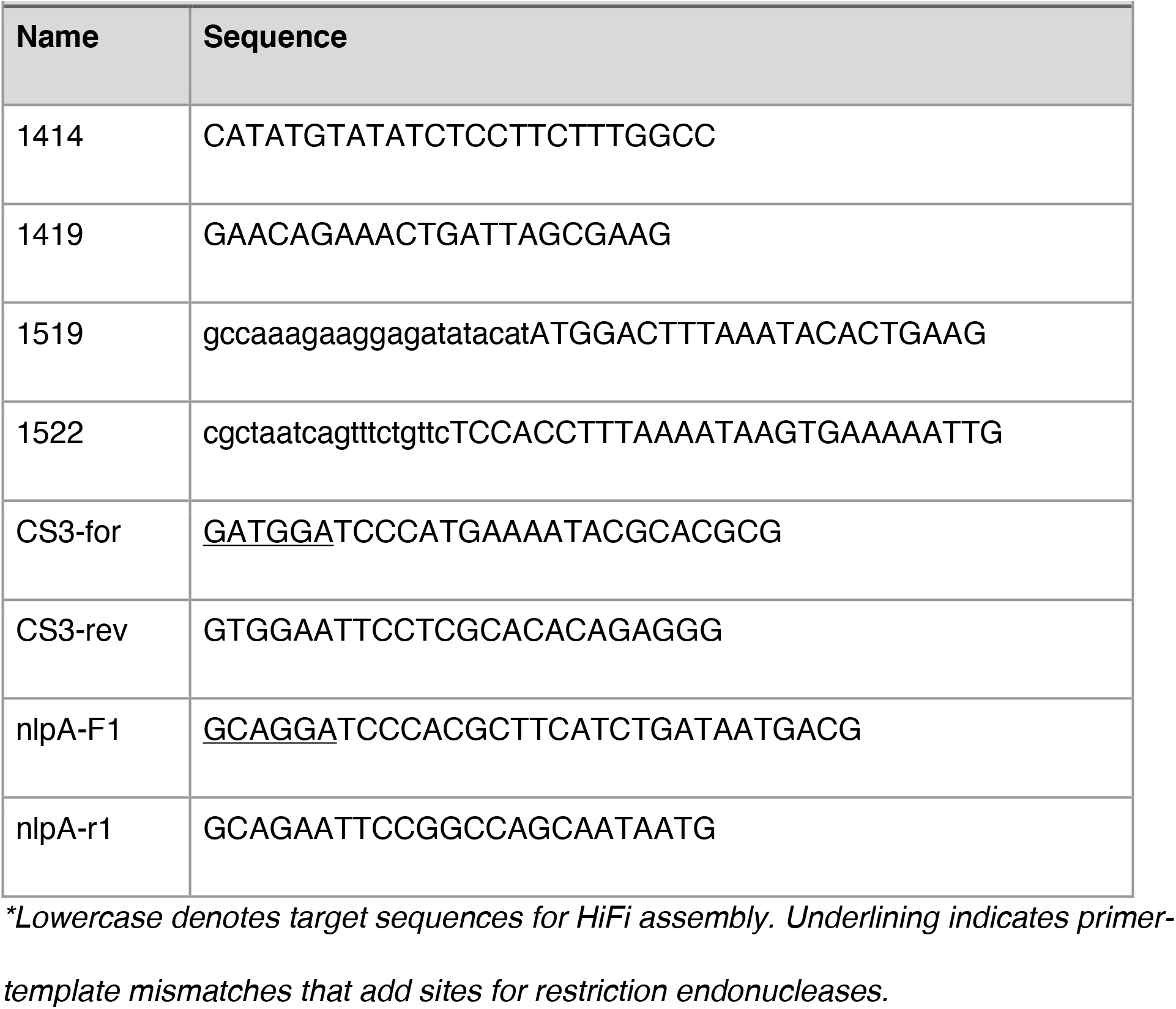
Oligonucleotides used in this study

### Reporter strains

Plasmid pHKLac1 is a promoterless *lacZ* reporter integration plasmid with a *pir*-dependent origin of replication (Haldimann and Wanner, 2001). It carries *attP*_HK022_ for Int_HK022_-mediated integration into the chromosome of *E. coli* at attB_HK022_. The *cs3* intergenic region (−121 to +352 relative to *cs3* ORF) was amplified from ETEC strain 1392-75-2a with primers CS3-for and CS3-rev. The *nlpA-yicS* intergenic region was amplified from ETEC strain H10407 with primers nlpA-F1 and nlpA-r1(Bodero *et al*., 2007). The PCR products were digested with BamHI and EcoRI and then ligated into the same sites of pHKLac1 to construct pCS3Lac1 [*cs3p*(−121 to +352 relative to ORF)*::lacZ*] and pNlpALac1 [*nlpAp*(−391 to +58 relative to ORF*)::lacZ*]. Each reporter plasmid was then integrated into the chromosome of MC4100 *[F-araD139 Δ(argF-lac)U169 rpsL150* (StrR*) relA1 flhD5301 deoC1 ptsF25 rbsR]* (Casadaban, 1976) as previously described (Haldimann and Wanner, 2001) resulting in strains GPM1072 (*attB*_HK022_::pCS3Lac1) and GPM1080 (*attB*_HK022_::pNlpALac1). Colony PCR was used to verify that each strain possessed only a single plasmid integrant, as previously described (Haldimann and Wanner, 2001). Strains used in this study are listed on table 5.

**Table 5:** Strains used in this study

| Strain | Characteristics | Reference |
| --- | --- | --- |
| MC4100 | *E. coli* K-12 *F*- *araD139* $\Delta$(*argF-lac*)*U169 rpsL150*<br>(StrR) *relA1 flhD5301 deoC1 ptsF25 rbsR* | (Casadaban, 1976) |
| GPM1072 | MC4100 *attB*<sub>HK022</sub>::pCS3Lac1 | This study |
| GPM1080 | MC4100 *attB*<sub>HK022</sub>::pNlpALac1 | (Bodero *et al.*, 2007) |
| H10407 | ETEC O78:H11 CFA/I+ ST+ LT+ CexEα+ | (Skerman *et al.*, 1972) |
| 1392/75-2a | ETEC O6:H16 CS1+ CS3+ ST+ LT+ CexEκ+ CexEε+ | (Adlerberth *et al.*, 1995) |

### B-galactosidase assays

Lac reporter strains GPM1072 and GPM1080 were transformed with pGPMRns-Myc or vector pTags2 to determine the effect of decanoic acid on the ability of Rns to regulate Rns-dependent promoters. All strains were grown aerobically at 37C to stationary phase in LB medium with 100µg/mL ampicillin with or without decanoic acid in 0.4% vol/vol DMSO. B-galactosidase activity was assayed as previously described (Miller, 1972).

### Immunoblots

ETEC strains H10407 and 1392/75-2a were grown to stationary phase in CFA (Skerman *et al*., 1972; Adlerberth *et al*., 1995) medium with or without decanoic acid in 0.4% vol/vol DMSO. Pilins were released from the outer membrane by incubating ETEC in 1/10 volume PBS at 65C for 20 minutes. Supernatants were clarified by centrifugation to remove insoluble material. Stationary phase whole cell lysates of ETEC or 10 µg pilin supernatants were subjected to SDS-PAGE, transferred to PVDF membranes, and blocked in TBS-Blotto (25 mM TrisCl pH 7.6, 150 mM NaCl, 5% wt/vol powdered nonfat milk. Antibodies against CexEα (H10407) and CexEε (1392/75-2a) were produced by immunization of rabbits (Proteintech Group, Inc.) with purified antigens and used at a dilution of 1:5,000 in TBS-Blotto with 0.05% vol/vol Tween20 (Pilonieta *et al*., 2007; Rivas *et al*., 2020) Primary antibodies against c-Myc (Sigma-Aldrich M4439) and DnaK (AbCam ab69617) were used at dilutions of 1:5,000 and 1:10,000 respectively. HRP conjugated goat anti-rabbit (Santa Cruz Biotechnology sc-2030) and goat anti-mouse (Jackson ImmunoResearch 115-036-062) antibodies were used at 1:10,000 dilutions. Chemiluminescence was detected with an Odyssey FC Imaging System (LI-COR Biosciences).

## Results

### Identification of proteins potentially regulated by UFAs

Secondary structure prediction and alignments identified eight AraC proteins that are known virulence regulators and that we predict may be regulated by UFAs (Fig 1, Table 1). Many of these proteins did not contain a lysine in the area corresponding to Lys230 in ToxT, but rather an arginine, and in one case histidine. When these proteins were compared to an existing phylogenic tree of fifty-eight AraC family proteins (Ibarra *et al*., 2008), all were found to be among the most closely related to ToxT. This supports our hypothesis that these AraC family members could share a common mechanism of being regulated by fatty acid ligands and served as a foundation for subsequent structural and functional analysis of these target proteins.

### Initial structure determination

As we pursued several of these targets for biochemical and structural studies, we were able to crystallize and obtain diffraction data for Rns. As attempts at molecular replacement using ToxT and other AraC family member structures as phasing models were unsuccessful, we used SeMet labeling coupled with single-wavelength anomalous dispersion (SAD) to experimentally determine phases and solve the structure. As expected, Rns shows the hallmarks of AraC proteins, with a cupin-like N-terminal domain and helix-turn-helix containing DNA binding domain (DBD) (Fig 2A). The N-terminal domain (residues 1-158) contained 6 β-strands and 4 α-helices. The fourth helix is connected through a three residue linker to the DBD. The DNA binding domain (residues 162-265) is typical for this family of proteins, consisting of seven alpha helices. Interestingly, one monomer was found to be in an open conformation in which the N- and C-terminal domains were separated, forming a groove containing electron density consistent with a bound ligand. In addition, the protein formed a dimer across a crystal contact in the N-terminal domain similar to other AraC proteins that are known to dimerize (Soisson *et al*., 1997; Shrestha *et al*., 2015).

**Figure 2:**
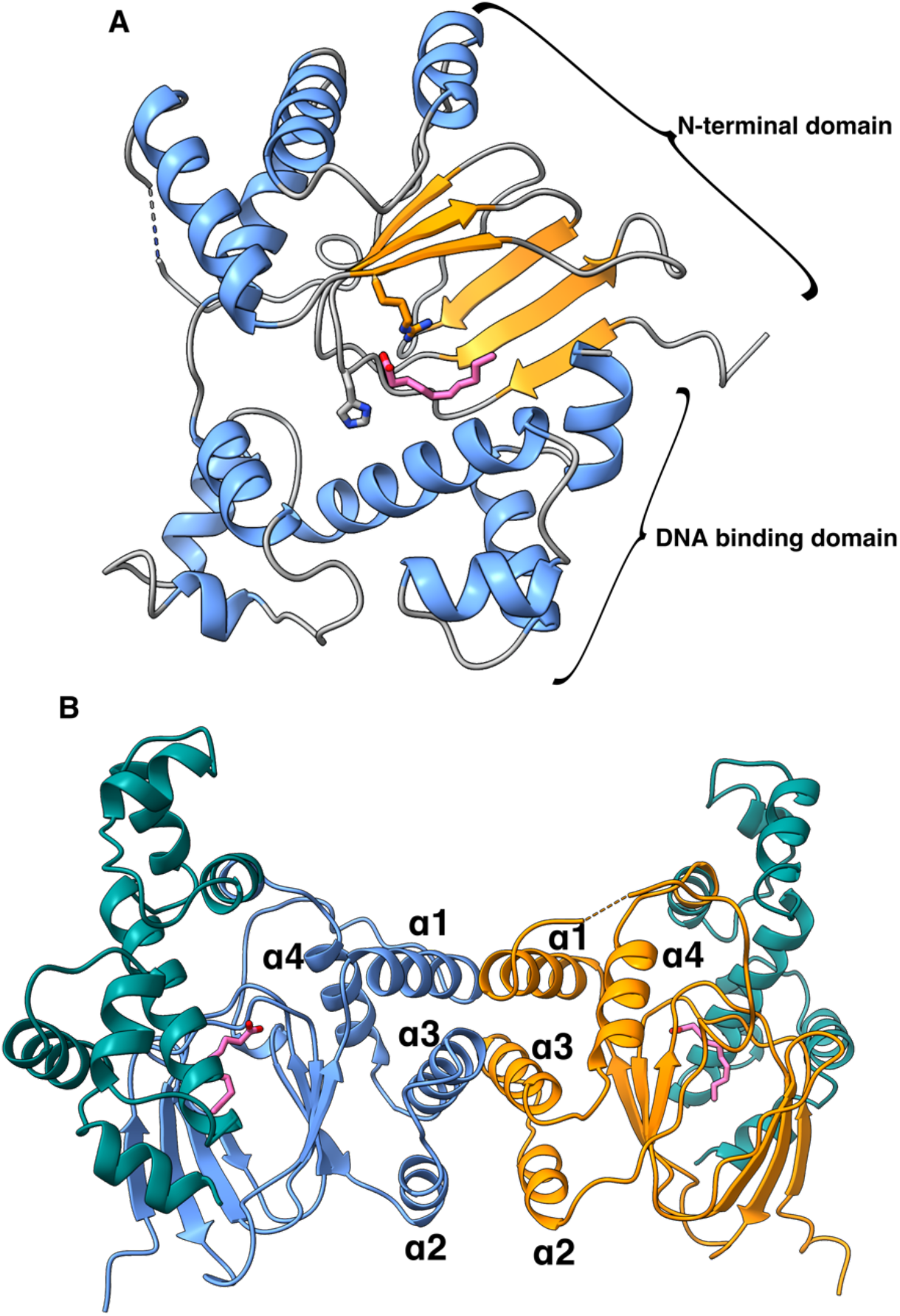
Structure of Rns and the potential dimerization interface. **A**. Structure of SeMet-Rns showing decanoic acid modeled in the ligand binding pocket with α-helices in blue, β-sheets in orange, coils in gray, and the decanoic acid in pink. The N-terminal and DNA binding domains are labeled. **B**. Proposed biological dimer with the N-terminal domains of the two monomers in orange or blue and both DNA binding domains in teal. The N-terminal α-helices are labeled.

### Rns has a dimer interface

Rns formed a dimer across a crystallographic symmetry axis with helices α1-α3 from the N-terminal domain forming an interface consistent with physiological function (Fig 2B). The helices from the two monomers face each other in an antiparallel fashion in a manner similar to ExsA, another AraC family member from *Pseudomonas aeruginosa* involved in the transcription of the type three secretion system (Frank and Iglewski, 1991; Shrestha *et al*., 2015). In Rns, much of the interface is between α1 and α3, burying approximately 274 Å^2^ of surface area. The interface forms in such a way that the DNA binding domains of the two Rns monomers are pointing almost 180° away from each other (Fig 2B), implying Rns is capable of binding looped DNA in a manner similar to AraC (Lobell and Schleif, 1990). Ongoing studies in our laboratories are focused on investigating the relevance of the dimerization interface.

### Rns contains an unexpected ligand

Analysis of the crystal structure revealed unexpected electron density in the groove between the N- and C-terminal domains consistent with the presence of a medium chain fatty acid. Initially, octanoic acid, an eight-carbon fatty acid was added to the model. While this improved the R factor, the density appeared to be able to accommodate a larger molecule.

When decanoic acid was modeled, the refinement statistics improved, and the two additional carbons filled the remaining electron density (Fig 3A). As no exogenous fatty acids were added to Rns during purification or crystallization, the presence of bound fatty acid in the structure suggested a possible physiological role, and therefore we proceeded with functional analysis of the effect of fatty acids on Rns activity.

**Figure 3:**
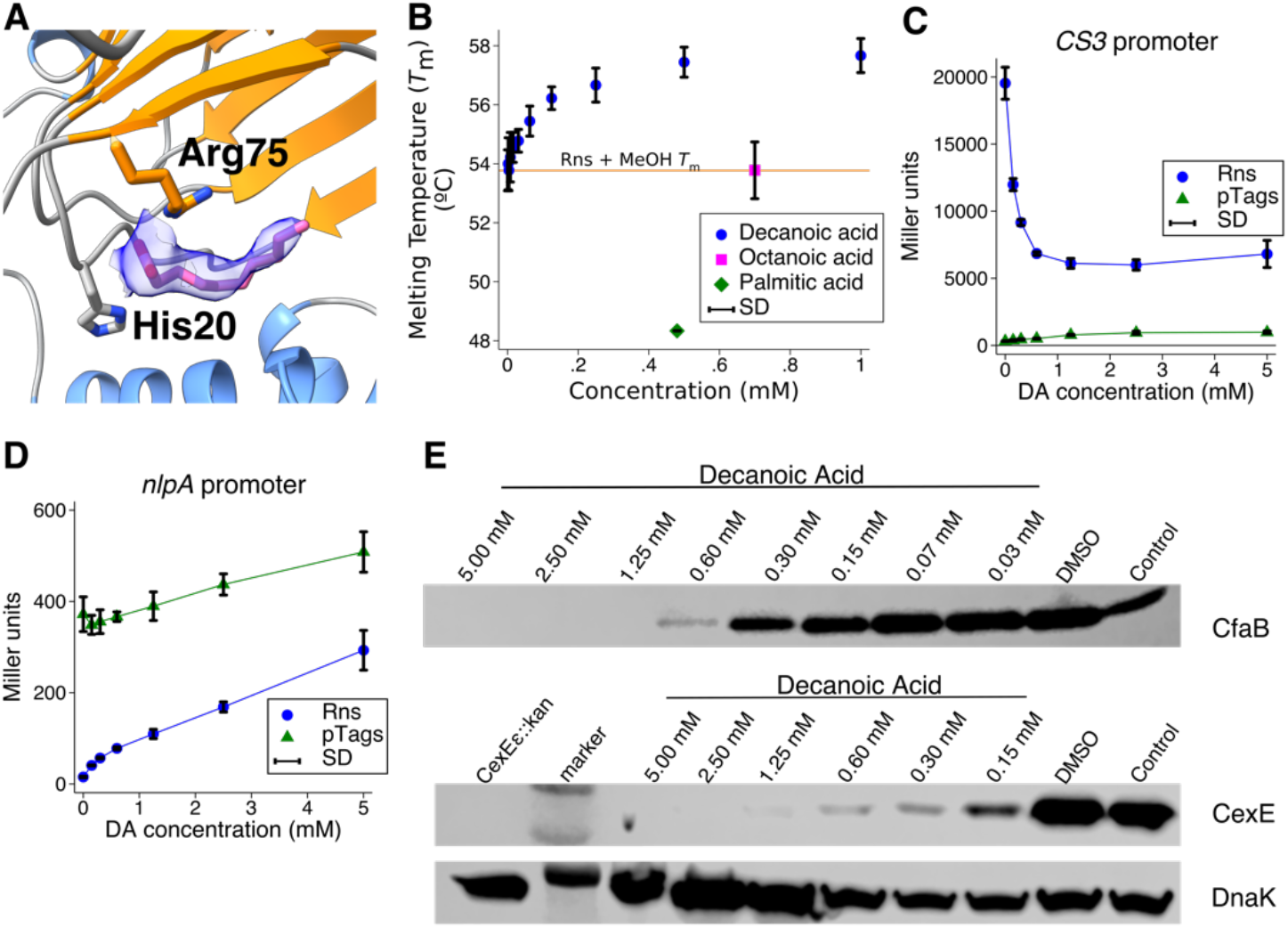
Rns binds decanoic acid, which influences Rns activity. **A**. Detail showing the electron density from an 2Fo-Fc omit map contoured to 1σ around the decanoic acid as a blue volume. **B**. DSF showing an increase in the *T*m of Rns in a dose dependent manner with decanoic acid. Octanoic acid does not affect the *T*m of Rns and palmitic acid decreases the melting temperature of Rns. Data is shown as the mean ± SD, n=3. **C**. B-galactosidase assays showing decanoic acid represses Rns activity at the *CS3* promoter and **D**. causes Rns derepression at the *nlpA* promoter. Data is given as the mean response ± SD, n=3. **E**. Western blots of CfaB (top) and CexE (lower) showing decreased expression when increasing concentrations of decanoic acid were added to the culture.

### Decanoic acid binds to Rns and inhibits its activity

Differential scanning fluorometry (DSF) was used to determine if decanoic acid and other fatty acids could interact with Rns. DSF experiments were performed with decanoic acid (10 carbons), octanoic acid (8 carbons), and palmitic acid (16 carbons). While octanoic acid did not change the melting temperature (*T*_m_) of Rns, palmitic acid decreased its *T*_m_ and decanoic acid increased the *T*_m_ in a dose dependent manner up to a concentration of 1 mM (Fig 3B). This is consistent with decanoic acid interacting specifically with Rns.

As the DSF showed decanoic acid interacts with Rns, we evaluated its effect at Rns activated (*CS3p*) and repressed (*nlpAp*) promoters using Lac reporter strains (Favre *et al*., 2006; Bodero *et al*., 2007). Consistent with the structure and DSF results, addition of exogenous decanoic acid inhibited Rns-dependent expression of β-galactosidase from *CS3p* in a dose-dependent manner (Fig 3C). Inhibition was not the result of nonspecific interactions, as exogenous decanoic acid also relieved Rns-dependent repression of the *nlpAp*, mirroring the effects at the *CS3p* (Fig 3D). As both repression and activation require Rns binding at sites near each promoter ((Bodero *et al*., 2007), and Munson, unpublished), these results suggest decanoic acid directly interferes with the ability of Rns to bind DNA.

To determine if decanoic acid can inhibit in situ virulence gene expression, the fatty acid was added to ETEC cultures. Consistent with the results from K-12 Lac reporter strains, we found that increasing concentrations of decanoic acid inhibited the expression of the CFA/I major pilin CfaB in ETEC H10407 (Fig 3E top panel). This was not limited to H10407 because the expression of CexEε from ETEC strain 1392/75-2a was also inhibited by the presence of decanoic acid (Fig 3E middle panel). The apparent lack of an effect upon DnaK expression suggest decanoic acid specifically inhibits the Rns regulon (Fig 3E bottom panel). Thus, exogenous decanoic acid abolishes the expression of ETEC virulence factors by inhibiting the activity of the Rns transcription factor. This is the first evidence of a small molecule that specifically inhibits Rns mediated ETEC virulence and has significant implications for therapeutic applications.

### Structure with decanoic acid

Given decanoic acid both stabilized and affected Rns function, Rns was crystalized in the presence of excess decanoic acid to assess what effects this might have on its structure. Despite the diffraction data showing significant pseudo-translation, the structure was solved to 3.0 Å using molecular replacement with the apo SeMet-Rns structure as a search model. Analysis of the resulting electron density showed both monomers contained a ligand, and decanoic acid was placed in the density.

Overall, the SeMet and the unlabeled Rns structures are quite similar. Interestingly, in the unlabeled structure both monomers were in a closed conformation. Therefore, we examined differences between the open monomer observed in the SeMet structure and a closed monomer of the unlabeled structure. In the SeMet structure, the ligand binding groove is open as a result of helices making up the dimer interface and β-strands 2, 4, and 6 shifting away from the long helix in the DBD by ∼1.7 Å in comparison to the native Rns structure (Fig 4A and B). This shift results in the binding pocket closing in around decanoic acid (compare Fig 4C to 4D).

**Figure 4:**
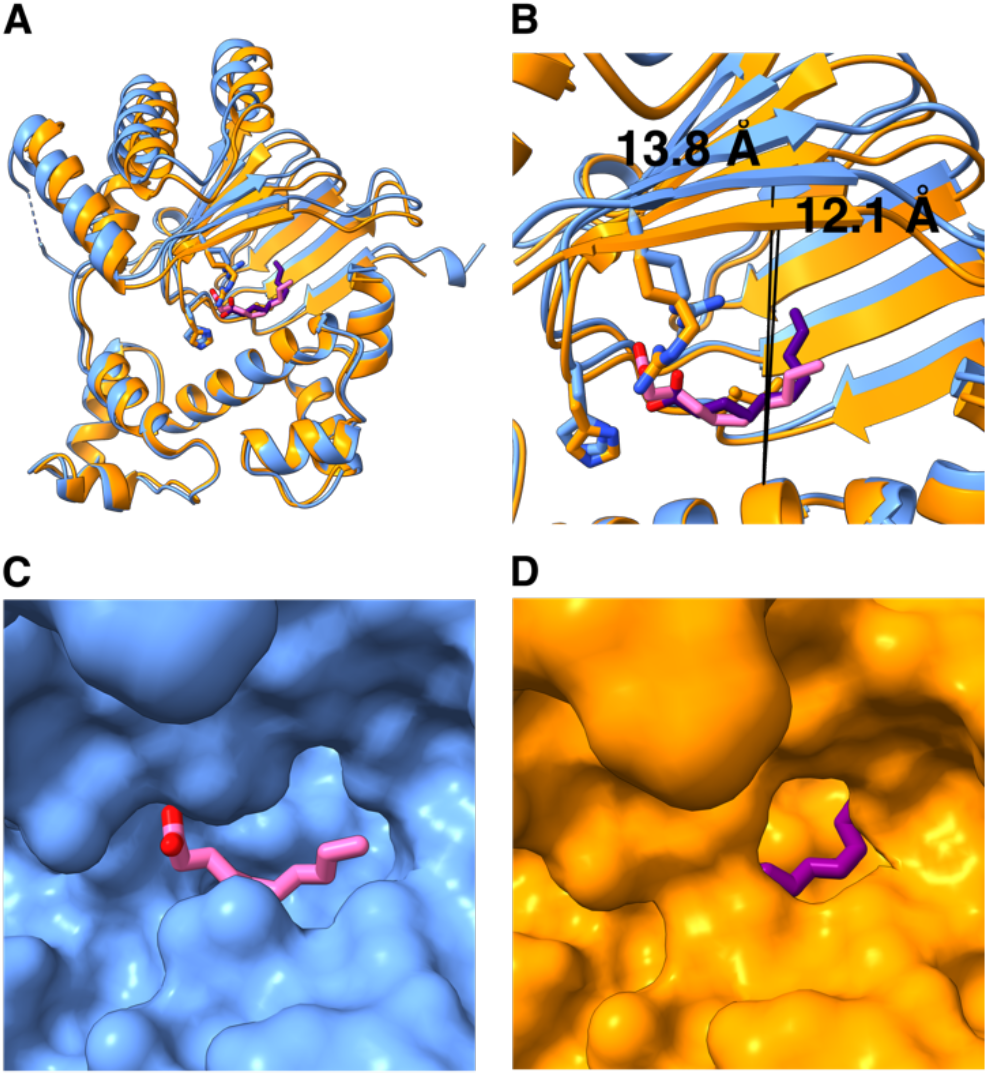
Comparison of the SeMet labeled structure in the open conformation and the native Rns structure in the closed conformation. **A**. Superposition of the native (orange) and SeMet-Rns (blue) structures, the two domains were aligned using the long helix in the DNA binding domain (Phe205-Glu223), showing how the N-terminal domain is shifted away from the DBD in the SeMet structure. **B**. Detail showing the β-sheet Is shifted by ∼2 Å from the DBD in the SeMet structure compared to the native structure. **C**. Surface representation of the SeMet-Rns structure in blue with the decanoic acid in pink showing how the fatty acid is exposed to solvent. **D**. Surface representation of the native-Rns in orange with the decanoic acid in purple showing how the ligand is enclosed in more of a pocket.

### Arg75 and His20 in the structures make contact with the decanoic acid

Both the open and closed conformations show His20 and Arg75 interacting with the carboxyl group of the decanoic acid (Fig 5A and 5B). We note that Lys216, which was the positively charged residue identified by our computational screen as potentially interacting with the fatty acid (Fig. 1), does not interact with the ligand.

**Figure 5:**
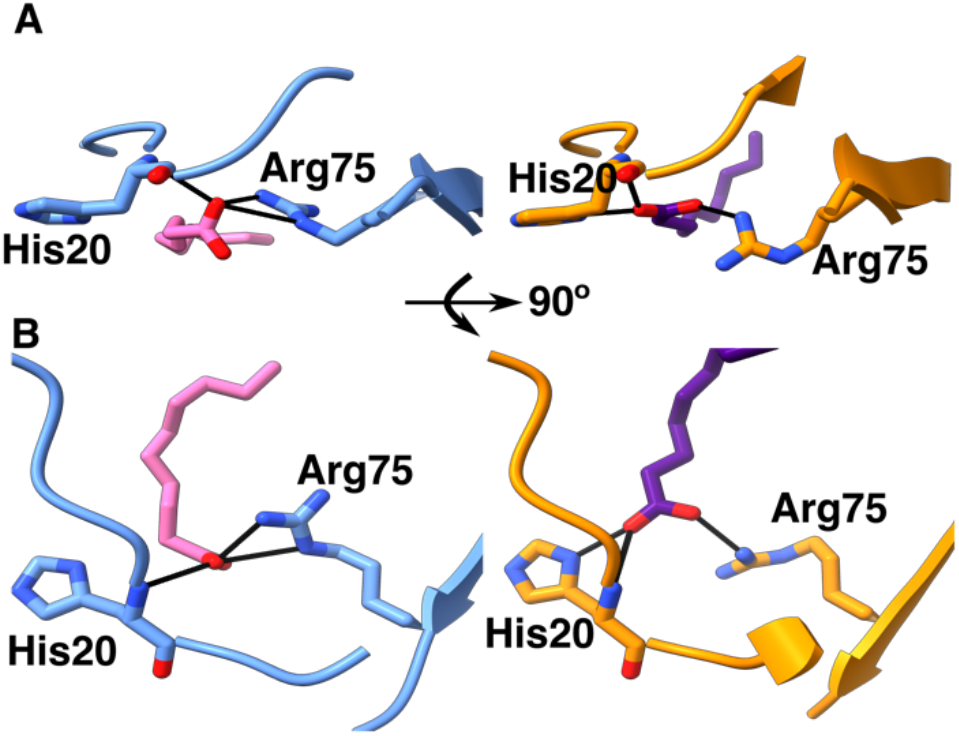
Detail showing the interactions between His20 and Arg75 to decanoic acid. **A**. “front” view showing how the Arg75 in the SeMet-Rns, in blue, is above the decanoic acid (left panel) while in the native-Rns, in orange, Arg75 (right panel) is alongside the fatty acid. **B**. “Top” view with the SeMet-Rns in blue (left panel) and native-Rns in orange (right panel).

Interestingly, in the open structure Arg75 is between the fatty acid and the rest of the N-terminal domain (Fig5A left panel), compared to the closed Rns where Arg75 is more alongside the ligand (Fig 5A right panel). In addition, the position of the decanoic acid head group is in a different orientation in the open and closed conformations. In the open conformer the head group is perpendicular to the plane of the imidazole group of His20 resulting in only one oxygen from the fatty acid making contacts with His20 and Arg75 (Fig5A and B left panel). In the closed conformation the fatty acid carboxyl group is turned 90° allowing both oxygens to make contacts with either His20 and Arg75 (Fig 5A and B right panel). It is interesting to speculate that the position of the Arg75 drives the transition between open and closed conformations. The role of Arg75 and His20 in mediating the structural response of Rns binding to decanoic acid is being explored.

### Comparison of Rns and ToxT binding pockets

Given Rns and ToxT are the only full-length AraC structures solved with bound fatty acids it is interesting to compare the fatty acid binding mode of the two proteins. Based on crystal structures, the proteins bind fatty acids of different lengths and degree of saturation: palmitoleate (16 carbon monounsaturated) binds to ToxT, while decanoate (10 carbon saturated) binds to Rns. To compare the ToxT structure (3GBG) (Lowden *et al*., 2010) to the native Rns structure, residues 211-231 from ToxT were aligned with residues 205-223 from Rns in Chimerax (Goddard *et al*., 2017). While both fatty acids are bound between the N-terminal domain and the DBD (Fig 6), in ToxT the bound fatty acid projects more deeply into the N-terminal domain (compare Fig 6A to 6B). Interestingly, in ToxT residues from the N-terminal domain (Lys31) and the DBD (Lys230) interact with carboxyl group on the fatty acid (Fig 6A), while in Rns the two residues that interact with the carboxyl group are in the N-terminal domain (Fig 6B). Furthermore, the β1-strand in ToxT is absent in the Rns structure (compare Fig 6A to 6B), creating a more enclosed pocket in ToxT (Fig 6C), whereas Rns has an opening in the side of the pocket (Fig 6D). Both pockets are hydrophobic except for the charged groups involved in binding the carboxyl portion of the fatty acid (Fig 6C and 6D). In addition, palmitoleate occupies a larger portion of the binding pocket in ToxT than the decanoate does in Rns (Fig 6C and 6D), suggesting Rns is able to bind to fatty acid ligands larger than decanoate. It is interesting to note while fatty acids regulate both proteins, the details of the binding are different and this is likely to be a theme across fatty acid regulated AraC’s.

**Figure 6:**
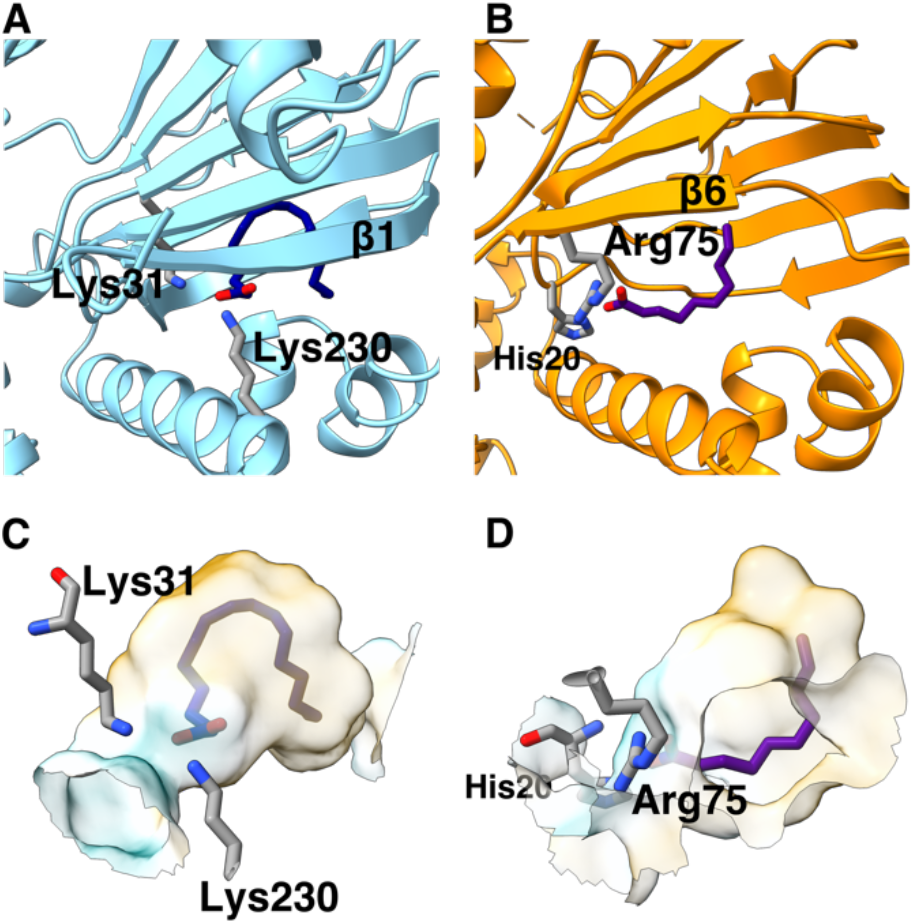
Comparison of the binding pockets of ToxT and Rns. **A**. ToxT in blue with the palmitoleic acid in dark blue. **B**. The native Rns in orange with the decanoic acid in purple were aligned in Chimerax (Goddard *et al*., 2017) using the long helix in the DBD. The β-strand for each protein closest to the long helix in the DBD are labeled, as well as the residues which make contacts with the fatty acids. The binding pockets for **C**. ToxT and **D**. Rns are shown. The surfaces are colored according to lipophilicity calculated using Chimerax (Goddard *et al*., 2017), with cyan for charged areas and orange for hydrophobic surfaces. Again, the residues interacting with fatty acids are labeled.

## Discussion

As part of ongoing efforts to determine whether other AraC proteins are regulated by fatty acids in a similar fashion as ToxT, we solved the structure of Rns using X-ray crystallography in two conformations: open, distinguished by having a groove between the N and C-terminal domains for ligand binding; and closed, where the two domains have come closer together with the pocket enclosing the ligand. In addition, the Rns structures were dimeric, with an interface similar to structures of other AraC proteins (Soisson *et al*., 1997; Shrestha *et al*., 2015). The DNA binding domains of the two monomers are pointing 180° apart from each other, suggesting Rns binds to DNA loops in a manner similar to AraC (Lobell and Schleif, 1990), and consistent with the observation that most Rns regulated promoters have a proximal site and a distal site separated by about forty base pairs, although at some promoters this is less and at others the sites are separated by over a hundred base pairs (Munson and Scott, 1999; Munson and Scott, 2000; Bodero *et al*., 2007; Pilonieta *et al*., 2007; Bodero *et al*., 2008; Bodero and Munson, 2016). For instance, *CS1* pili expression is mediated by two sites, one proximal to the −35 RNAP binding site, and one distal, around −106, to the transcription start site (Munson and Scott, 1999). The sites are asymmetric and contribute additively to the expression of the *CS1* pili. Another example is the *rns* promoter has two Rns binding sites required for transcription, one at the −227 position and the other downstream of the transcription start site (Munson and Scott, 2000). In this instance the binding sites are widely spaced and act synergistically as both binding sites are required for activating transcription (Munson and Scott, 2000). These findings, coupled with the observations from our structures, indicate Rns likely binds to looped DNA. Work is ongoing to investigate the relevance of this interface and to clarify if Rns binds looped DNA.

Importantly, the ligand binding pocket of Rns was occupied by the 10-carbon fatty acid, decanoic acid. DSF studies showed decanoic acid specifically stabilized Rns, as octanoic and palmitic acids did not. Decanoic acid also influenced Rns activity at the *CS3* promoter, where Rns activates transcription, and at the *nlpA* promoter, where Rns represses transcription (Bodero *et al*., 2007; Bodero *et al*., 2008). In addition, decanoic acid was able to repress the expression of Rns regulated CexE and the CFA/I pilin in ETEC strains. Furthermore, structural analysis has identified possible residues involved in binding to decanoic acid. Therefore, we have identified a small molecule that binds to Rns inhibiting ETEC virulence expression.

While the specific structural mechanism by which Rns is regulated by decanoic acid is not clear, the mechanism by which fatty acid inhibition occurs is likely linked to influencing the stability of the Rns dimer, either directly or by a dynamic allosteric mechanism as demonstrated for *V. cholerae* ToxT (Cruite *et al*., 2019). Future studies will be directed at clarifying this mechanism. The inhibition of Rns by a fatty acid represents the first time a small molecule has been shown to bind to Rns and inhibit virulence gene expression in ETEC. Decanoic acid and its analogs should be pursued as lead anti-virulence compounds, as they could be viable alternatives to decades of failed strategies to develop an effective vaccine against ETEC.

## Author Contributions

Charles R Midgett Roles: Writing-Original Draft Preparation, Formal Analysis, Visualization, Investigation, Project Administration; Kacey M Talbot Roles: Writing-Review and Editing, Investigation, Formal Analysis, Visualization; Jessica L Day Roles: Investigation; George P Munson Roles: Writing-Review and Editing, Funding Acquisition, Project Administration, Methodology, Project Administration; F Jon Kull Roles: Writing-Review and Editing, Funding Acquisition, Project Administration, Conceptualization, Project Administration

## Data Availability

The structures of Rns have been deposited into the PDB: SeMet-Rns; 6XIV, and Rns; 6XIU.

## Acknowledgments

Research reported in this publication was supported by the NIAID of the National Institutes of Health under award number AI128164 (GPM) as well as AI072661 and AI140740 (FJK). Additional support was provided by BioMT COBRE P20-GM113132. Sequencing was performed by the Molecular Biology Shared Resource at Dartmouth. This research used the FMX and AMX beamlines of the National Synchrotron Light Source II, a U.S. Department of Energy (DOE) Office of Science User Facility operated for the DOE Office of Science by Brookhaven National Laboratory under Contract No. DE-SC0012704. The Life Science Biomedical Technology Research resource is primarily supported by the National Institute of Health, National Institute of General Medical Sciences (NIGMS) through a Center Core P30 Grant (P30GM133893), and by the DOE Office of Biological and Environmental Research (KP1605010). We also acknowledge the beam line staff at NSLS2-FMX and AMX for making data collection possible. The contents of this publication are solely the responsibility of the authors and do not necessarily represent the official views of the National Institutes of Health.

